# Aging modulates the effects of scene complexity on visual search and target selection in virtual environments

**DOI:** 10.1101/2023.05.18.540524

**Authors:** Isaiah J. Lachica, Aniruddha Kalkar, James M. Finley

## Abstract

Processing task-relevant visual information is important for many everyday tasks. Prior work demonstrated that older adults are more susceptible to distraction by salient task-irrelevant stimuli, leading to less efficient visual search. However, these studies often used simple stimuli, and less is known about how aging influences visual attention in environments more representative of real-world complexity. Here, we test the hypothesis that aging impacts how the visual complexity of the environment influences visual search. Young and older adults completed a virtual reality-based visual search task in environments with increasing visual complexity. As visual complexity increased, all participants exhibited longer times to complete the task which resulted from increased time transferring gaze from one correct target to the next and increased delay between when correct targets were fixated and selected. The increase in time to completion can also be attributed to longer times spent re-fixating task-relevant objects and fixating task-irrelevant objects. These changes in visual search and target selection with increasing visual complexity were greater in older adults and working memory capacity was associated with multiple performance measures in the visual search task. These findings suggest that visual search performance could be integrated into assessments of working memory in dynamic environments.

## Introduction

Selecting and processing relevant visual information from the environment is vital for planning and executing everyday tasks (1–4). For example, when walking in environments where precision stepping is required, we use vision to plan routes, step over obstacles, or step onto safe footholds (5–7). However, the ability to properly direct vision to relevant visual stimuli appears to be impaired in older adults. While walking, older adults appear to be more easily distracted by future threats on their path than young adults as they shift their gaze away from the current stepping target to the next before completing the step (8,9). This gaze behavior can be problematic as it can result in greater stepping errors (10). Understanding how we select and process visual information in the environment and how these processes change with aging is crucial for preventing potentially injurious behaviors.

Natural and human-made environments are filled with visual stimuli that may be relevant or irrelevant for guiding future actions. Which visual stimuli are selected and processed may depend on how overt visual attention is directed. The control of visual attention has been prominently described as a balance between bottom-up and top-down processes. Bottom-up processes emphasize saliency, a characteristic of visual stimuli based on low-level visual features such as contrast and luminance (11), which makes highly salient stimuli “pop out” of a scene and capture attention automatically. In contrast, top-down processes control visual attention based on cognitive processes that prioritize factors such as task demands, prior knowledge, or gist of realistic scenes (12–15). Task-dependent trade-offs between these processes enable attention to be directed based on top-down factors when performing a task or by highly salient stimuli when necessary. However, the balance between bottom-up and top-down control appears to rely on an individual’s cognitive capacity, which is affected by normal aging (16,17).

Older adults appear more susceptible to distraction by irrelevant visual stimuli than young adults due to age-related changes in their cognition, particularly in their ability to exert top-down inhibition (18–20). These effects prevent older adults from suppressing the automatic capture of attention by salient task-irrelevant stimuli, negatively affecting their performance in visually demanding tasks (21–24). Impaired capacity for inhibition can also lead to task-irrelevant information being encoded into working memory at the cost of task-relevant information (22,25). However, prior studies on age-related changes in visual search used simple visual scenes composed of arbitrarily selected shapes and letters. It is unclear whether these age-related effects on visual attention extend to performing tasks in scenes more closely reflecting the visual complexity in everyday visual search.

The Trail Making Test (26) is a visual search task commonly used to assess cognitive domains such as visual attention, working memory, and inhibition (27–29). It is typically implemented in a pen-and-paper format with numbers and letters as search targets and completion time as the primary outcome measure. Performance in the test appearing to decline with age as older adults tend to take more time to complete the task (30–34). Recent attempts have been made to improve the generalizability of the test by transforming it into a three-dimensional pointing task in virtual reality (VR) (35). Studies have also supplemented completion time with eye-tracking measures, such as average fixation time and number of fixations, to improve the interpretability of test results and identify differences in experimental (36) and neurological (37,38) conditions. However, the context in which people search for targets in both test versions does not reflect the visual complexity of search targets or the level of distraction present in everyday visual search. Questions remain on how increasing visual complexity influences visual search in young and older adults.

We aimed to determine how aging-related differences in cognition influence visual search and target selection in increasingly complex virtual environments. Participants completed a custom-designed VR-based visual search task based on the Trail Making Test-B in three levels of increasing visual complexity (39). To determine the influence of visual attention on task performance, we recorded eye movements from which we calculated the time between fixations on correct targets, average and total fixation times, number of fixations, and the saliency of fixated regions. We also determined how the actions involved in target selection using a laser pointer influenced task completion time by measuring the number of selection errors and the delay between fixating and selecting a correct target. Additionally, all participants completed assessments of global cognition, short-term memory, working memory, and inhibitory capacity. We hypothesized that as the complexity of the visual scene increased, participants would take longer to complete the task because attention would be attracted by task-irrelevant stimuli, leading to a reduced capacity to encode correct targets in working memory. Specifically, all participants would be more prone to re-fixating search targets and fixating on salient, task-irrelevant distractors, leading to longer times between fixations on correct targets. In addition, we hypothesized that this performance decline would be greater in older adults due to aging-related impairments in their cognition.

## Methods

### Participants

15 young (9 female, age: 27.7 ± 3.33 years) and 15 older adults (9 female, age: 71.8 ± 4.46 years) with normal or corrected-to-normal vision participated in the study. Data from two older adults were excluded due to hardware/software issues. We conducted a sample size calculation for mixed-effects regression models using GLIMMPSE (40) and used pilot data to estimate the variance in our dependent variables (task completion time, total task-relevant fixation time, and saliency of fixated regions). We found the largest sample size estimate across our three dependent variables and determined that a sample size of 26 (13 for each age group) was needed for a target power of 0.80, assuming a medium effect size of 0.45. All study procedures were reviewed and approved by the University of Southern California’s Institutional Review Board. All participants provided written informed consent before participating and were provided monetary compensation for their time. All aspects of the study conformed to the principles described in the Declaration of Helsinki.

### Experimental protocol

All participants completed the paper-based Trail Making Test-B (26) to allow for comparison with their performance in the VR-based visual search task. Additionally, the participants completed a battery of tests selected to assess different domains of cognition including the Montreal Cognitive Assessment (41) for global cognition, the Corsi task (42) for short-term memory, the Backwards Corsi task (43) for working memory, and the Flanker (44) task for the ability to inhibit the direction of attention to task-irrelevant stimuli. The Montreal Cognitive Assessment was administered on paper while the rest of the cognitive assessments were completed on a computer using Psytoolkit (45,46).

After completing the cognitive assessments, the participants completed three familiarization trials of the visual search task. These trials were completed in virtual environments with only the search targets present in front of a blank background to emphasize the identity of the search targets. In the first familiarization trial, the experimenter guided the participant through a single repetition of the task to provide a clear description of the task goal, the visual and auditory feedback that they will encounter, and the search target sequence. The guided familiarization trial was followed by two trials where participants completed the task independently and were only provided feedback by the experimenter at the end. After the familiarization trials, the participants completed 30 trials of the visual search task presented in three sets of 10 trials, each corresponding to one of three visual complexity levels. Only the data from the last five trials at each visual complexity level were processed and analyzed to account for potential learning effects.

### VR task

All participants completed a VR visual search task (Fig. 1a) which we based on the principles of the Trail Making Test-B. We particularly incorporated the visual attention, working memory, and inhibition (27–29) components of the Trail Making Test-B into our VR task. Participants were instructed to search for and select targets alternating between letters of the alphabet and animals whose names start with those letters in ascending order (A, Ant, B, Butterfly, C, Chickens, D, Dog, E, Eagle, F, and Frogs). The search targets were positioned randomly for each trial in regions of the virtual environments where they would not be overly distinct from the background, such as across the sky for the letters and at ground level for the objects. The participants viewed the virtual environments using the HTC Vive Pro Eye (HTC, New Taipei, Taiwan) head-mounted display and selected targets using an HTC Vive Controller. The head-mounted display had a 110-degree field of view with a resolution of 1440×1600 pixels per eye (2880×1600 pixels combined). A laser pointer extended from the top of the controller in the virtual environment, and participants selected targets by aiming the laser at the center of the target and pulling the trigger on the controller. All participants were informed that they could freely move their heads in all directions. Visual and auditory feedback were provided to indicate whether the selected target was correct. Specifically, the participants heard a bell sound and saw a “Correct!” text appear in the environment when the correct target was selected. In contrast, the participants heard a buzzer sound and saw a “Try Again!” text appear when they selected an incorrect target or when they did not select the correct target properly (laser not aimed at the center of the target).

**Figure 1.**
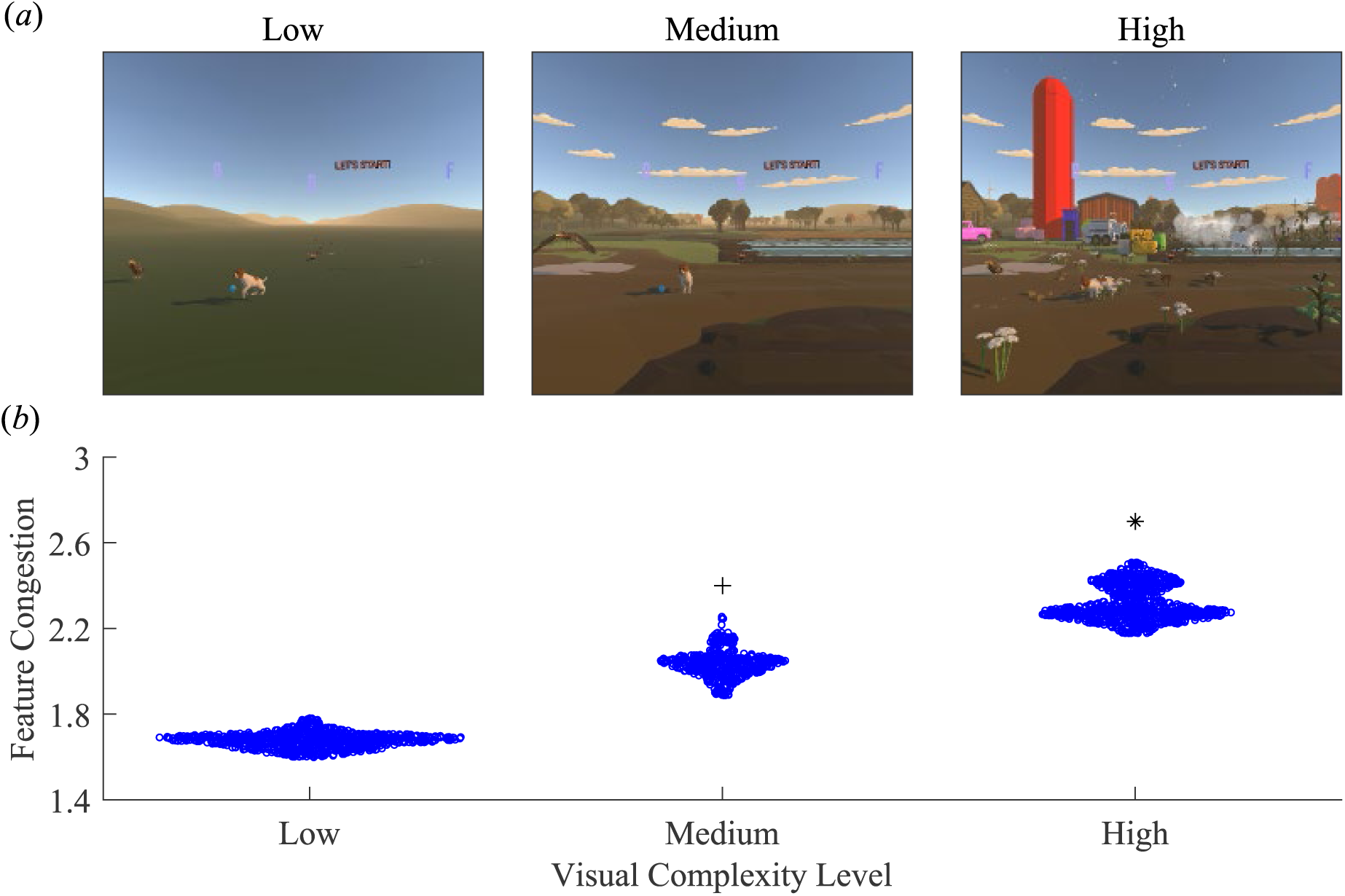
VR visual search task. (*a*) Single frames from representative gameplay recordings of each visual complexity level. Visual complexity was increased by adding visual distractors in the foreground and background in the form of vehicles, structures, plants, and agriculture-related inanimate objects to dissociate them from the search targets, which were the only letters and animals in the environment. (*b*) Swarm plot comparing feature congestion of each visual complexity level with each data point representing a frame from representative gameplay recordings. *Indicates significant difference between High and Low visual complexity levels and High and Medium visual complexity levels. ^+^Indicates significant difference between Medium and Low visual complexity levels.

The participants completed the visual search task in virtual environments with three levels of increasing visual complexity (Fig. 1a). The sequence of complexity levels was pseudo-randomized such that five participants in both young and older adult groups started with the low visual complexity level, another five with the medium visual complexity level, and the remaining five with the high visual complexity level. The visual complexity of the environment was manipulated by adding visual distractors in the foreground and background. These visual distractors came strictly in the form of vehicles, structures, plants, and agriculture-related inanimate objects to dissociate them from the search targets, which were the only letters and animals in the environment. Visual complexity was measured using feature congestion (47), which quantifies the distribution of low-level visual features of an image, including color, edge orientation, and luminance, as a single scalar measure with higher values indicating a more complex image. To determine if the modifications to the virtual environments produced increasing levels of visual complexity, we applied the feature congestion method to every frame of a representative gameplay recording from each visual complexity level (using Rosenholtz et al.’s Matlab implementation: https://dspace.mit.edu/handle/1721.1/37593). Feature congestion increased monotonically across visual complexity levels (Fig. 1b; *F*(2,2349) = 22550, *p* < 0.001). Post-hoc tests indicate that the high visual complexity level had greater feature congestion than the medium (Bonferroni-corrected *p* < 0.001) and low (Bonferroni-corrected *p* < 0.001) visual complexity levels. In addition, feature congestion was higher in the medium visual complexity level than in the low (Bonferroni-corrected *p* < 0.001).

### Data collection and processing

We collected the timing of each trigger press as participants selected objects. To quantify each participant’s overall performance, we calculated task completion time, which is defined as the difference between the time when the trial started and when the last correct target was selected. This measure of overall task performance is equivalent to the primary outcome measure of the paper-based versions of the Trail Making Test. Task completion time was calculated for every trial and reported as an average of the last five trials at each visual complexity level for each participant.

We collected eye and head movement data at 90 Hz using eye trackers and an inertial measurement unit built into the HTC Vive Pro Eye. The eye trackers were calibrated following a 5-point calibration process provided by HTC. Additionally, video recordings of the participants’ first-person point of view were recorded at 60 Hz for each trial using NVIDIA Shadowplay (NVIDIA, Santa Clara, CA, USA). All data processing was performed using a custom script in MATLAB R2022a (Mathworks, Natick, MA, USA). Horizontal and vertical eye-in-head angles were calculated from the unfiltered raw eye movement data and were combined with the horizontal and vertical rotations of the head to compute gaze angles. The gaze angles were then time-synchronized with the video recordings. Since the video recordings and the gaze angles did not have the same sampling rate, the gaze angles were then interpolated at each video recording frame.

As visual processing primarily occurs during periods of fixation(48,49), it was important to identify which samples qualified as such to determine how visual attention was deployed. To do so, we first computed horizontal and vertical gaze angular velocities by differentiating the interpolated gaze angles with respect to the time between video frames. We then identified fixations using a combination of a simple velocity threshold and a minimum fixation duration threshold. First, we categorized samples as fixations if their horizontal or vertical gaze angular velocity was less than 100 degrees per second(50). We computed gaze vectors for these fixations in the virtual environment with their origins set as the head position in the virtual environment and their direction determined by the gaze angles. We then recorded the names and locations of the objects or regions that the gaze vectors intersected in the virtual environment. We collapsed consecutive fixations to the same object or region together, calculated their durations, and removed fixations that were less than 60 milliseconds(51). We then classified a fixation as task-relevant if they were directed toward one of the search targets and task-irrelevant if they were directed toward the visual distractors or any other region in the environment. We further classified task-relevant fixations as either the first fixation on a search target in a trial or a re-fixation, which we define as fixations on search targets that were previously fixated.

We computed several measures of gaze-related task performance to capture how overt visual attention varied across conditions and between groups. We calculated the average time between fixations on correct targets by recording the time intervals from the end of a fixation on one correct target to the beginning of a fixation on the next correct target and then averaging these times within a trial. The average time between fixations on correct targets includes the saccade times between targets and any fixation time on incorrect targets. We also recorded the number of fixations on task-irrelevant objects and the number of re-fixations on task-relevant objects. Additionally, we calculated the average time spent on first fixations on task-relevant objects, re-fixations on task-relevant objects, and fixations on task-irrelevant objects by recording the time intervals from the start of a fixation on an object to the end of a fixation on the same object and averaging these times within a trial by fixation category. We similarly calculated the total time spent on first fixations on task-relevant objects, re-fixations on task-relevant objects, and fixations on task-irrelevant objects by recording the time intervals from the start of a fixation on an object to the end of fixation on the same object and finding the sum of these times within a trial by fixation category. These gaze-related task performance measures were calculated for every trial and reported as an average of the last five trials at each visual complexity level for each participant.

We generated saliency maps from each frame of the video recordings of the participants’ first-person point of view during the task using the original Itti, Koch, and Neibur(52) algorithm as implemented by Harel (J. Harel, A Saliency Implementation in MATLAB: http://vision.caltech.edu/∼harel/share/gbvs.php). Specifically, each video frame was decomposed into feature maps of its low-level visual features, which included color, orientation, and intensity. These feature maps were combined into a saliency map with corresponding scalar values for each pixel. These scalar saliency values were then converted to percentile ranks(53,54), with 100% indicating the most salient pixel and 0% the least salient pixel in the video frame. We recorded the percentile rank of each fixated region and then calculated saliency by averaging these recorded percentile ranks within a trial. We calculated saliency for every trial and reported it as an average of the last five trials at each visual complexity level for each participant.

We also recorded information on the objects selected at every trigger press, which included the identity of the selected object, the location where the laser pointer was being aimed in the virtual environment at the time of the trigger press, and whether the selection was correct. We then calculated from the number of selection errors, defined as selecting an incorrect target or misaiming (controller aimed outside the bounds of the correct target) when selecting a correct target, and selection delay, which is the time interval between fixating a correct target and successfully selecting it averaged within a trial. The number of selection errors and selection delay were calculated for every trial and reported as an average of the last five trials at each visual complexity level for each participant.

### Statistical analysis

All statistical analyses were performed in R (R Project for Statistical Computing) with the alpha value set at *p* < 0.05. Tests for normality and equal variances were performed using the Shapiro-Wilk test and Levene’s test with functions from the *stats* and *car* packages, respectively. Welch’s analysis of variance and pairwise Wilcoxon rank sum tests were used to compare the feature congestion of each visual complexity level using the *stats* package when assumptions of equal variance and normality were violated.

We fit linear mixed-effects models using the *lme4* package to test for the effects of age group, visual complexity, and their interaction on the following dependent variables: task completion time, average time between fixations on correct targets, average and total first fixation time on task-relevant objects, number of re-fixations on task-relevant objects, average and total re-fixation time on task-relevant objects, number of fixations on task-irrelevant objects, average and total fixation time on task-irrelevant objects, saliency, number of selection errors, and selection delay. We used the *lmerTest* package, which uses Satterthwaite approximations for the degrees of freedom, to test the null hypothesis that our model coefficients were zero. All models included random intercepts for each participant to account for repeated measures. Lastly, we performed post-hoc Bonferroni-corrected pairwise comparisons using the *emmeans* package when significant main effects or interactions, particularly differences between age groups within visual complexity levels, were found.

We compared cognitive assessment scores between age groups to determine the effect of age on various domains of cognition using functions from the *stats* package. Assumptions of normality and equal variance were tested using the Shapiro-Wilk and Levene’s tests, respectively. The Wilcoxon rank sum test was used when the normality assumption was violated; otherwise, a two-sample t-test was used.

To determine if our VR visual search task was able to mimic the paper-based Trail Making Test-B, we fit a multiple linear regression model using the *stats* package to identify any relationships between performances in both tasks. Specifically, we used Trail Making Test-B completion time as the response and VR task completion time and its interaction with visual complexity level as the predictors.

We fit three multiple linear regression models to test for associations between performance in the task and various cognitive domains. In our first model, we used VR task completion time in the high visual complexity level as our response and scores on the Montreal Cognitive Assessment, Corsi Block task, Backwards Corsi Block task, and the Flanker task as predictors. In our second model, we used the number of re-fixations on task-relevant objects as our response and scores on the Corsi Block task and the Backwards Corsi Block task as predictors. Finally, we used the number of fixations on task-irrelevant objects as our response and the Flanker task score as the predictor in our third model. We used the performance metrics recorded from the high visual complexity level as the range of values observed at this level covered those found at the lower visual complexity levels. We also included age as a predictor in all three models to control for age-related deficits in performance.

## Results

### Task completion times

All participants took longer to complete the task as the visual complexity of the environment increased, and this effect was greatest in older adults (Fig. 2). We found main effects of age group (*F*(1,26) = 49.19, *p* < 0.001) and visual complexity level (*F*(2,52) = 83.36, *p* < 0.001) on task completion time. We also found an interaction between age group and visual complexity level (*F*(2,52) = 9.50, *p* < 0.001). Compared to the young adults, the older adults took 8.9 seconds longer in the low complexity level (*p* = 0.002), 10.7 seconds longer in the medium complexity level (*p* < 0.001), and 17.7 seconds longer in the high complexity level (*p* < 0.001) to complete the task. In addition, the young adults took 8.2 seconds longer in the high versus low visual complexity (*p* < 0.001) and 7.5 seconds longer in the high versus medium visual complexity (*p* < 0.001), while the older adults took 17.0 seconds longer in the high versus low visual complexity (*p* < 0.001) and 14.4 seconds longer in the medium versus high visual complexity (*p* < 0.001) to complete the task.

**Figure 2.**
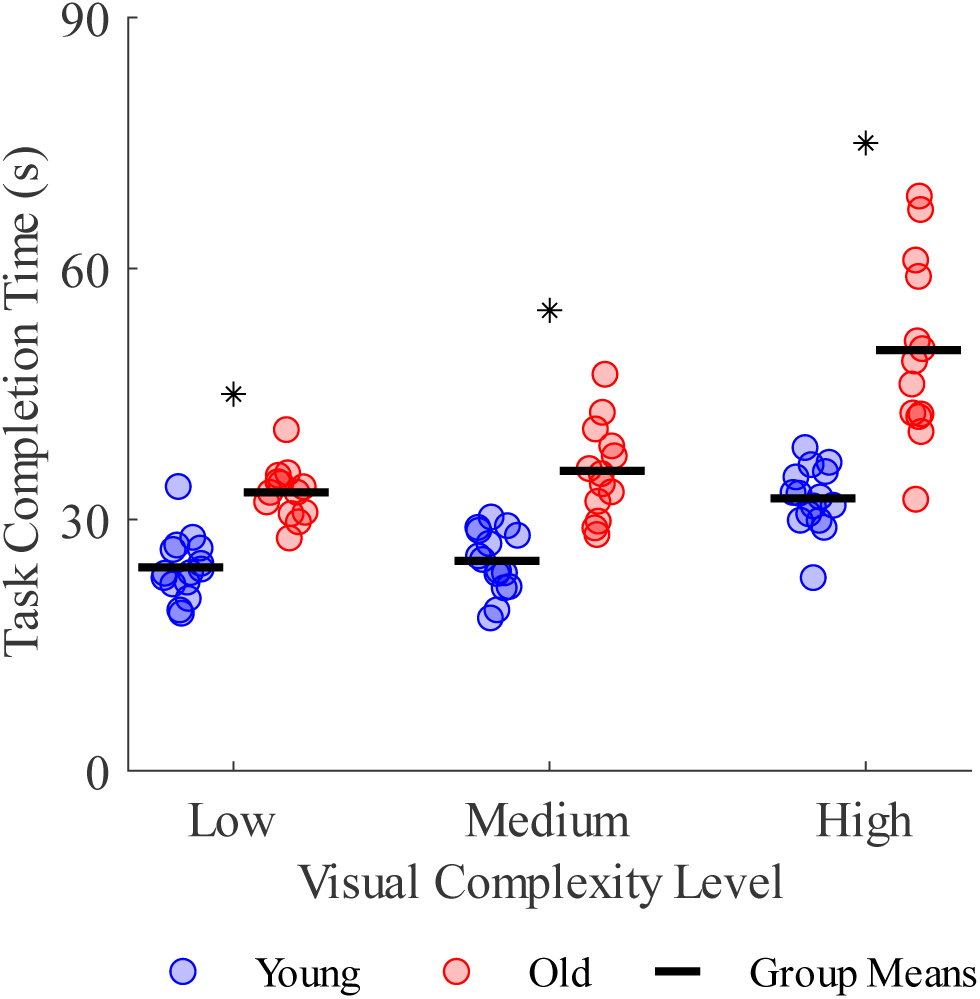
Effect of age group and visual complexity on VR task completion times. *Indicates significant difference between age groups.

### Time between fixations on correct targets

Several performance-related factors could explain the changes in task completion time with complexity level and differences in completion time between groups. For example, participants may have taken longer to complete the task in levels with higher visual complexity because they spent more time searching for correct targets. We calculated the average time between fixations on correct targets as the time interval from the end of a fixation on one correct target to the beginning of a fixation on the next correct target and found main effects of age group (*F*(1,26) = 42.13, *p* < 0.001) and visual complexity (*F*(2,52) = 83.69, *p* < 0.001; Fig 3a). We also found an interaction between age group and visual complexity (*F*(2,52) = 10.03, *p* < 0.001). Compared to young adults, older adults spent 0.6 seconds longer in the medium complexity level (*p* < 0.001) and 1.0 seconds longer in the high visual complexity level (*p* < 0.001) between fixations on correct targets. The effect of visual complexity level was larger in older adults, with young adults spending 0.5 seconds longer in the high versus low complexity level (*p* < 0.001) and in the high versus medium complexity level (*p* < 0.001), while the older adults spent 1.0 seconds longer in the high versus low complexity level (*p* < 0.001) and in the high versus medium complexity level (*p* < 0.001) on average between fixations on correct targets.

**Figure 2.**
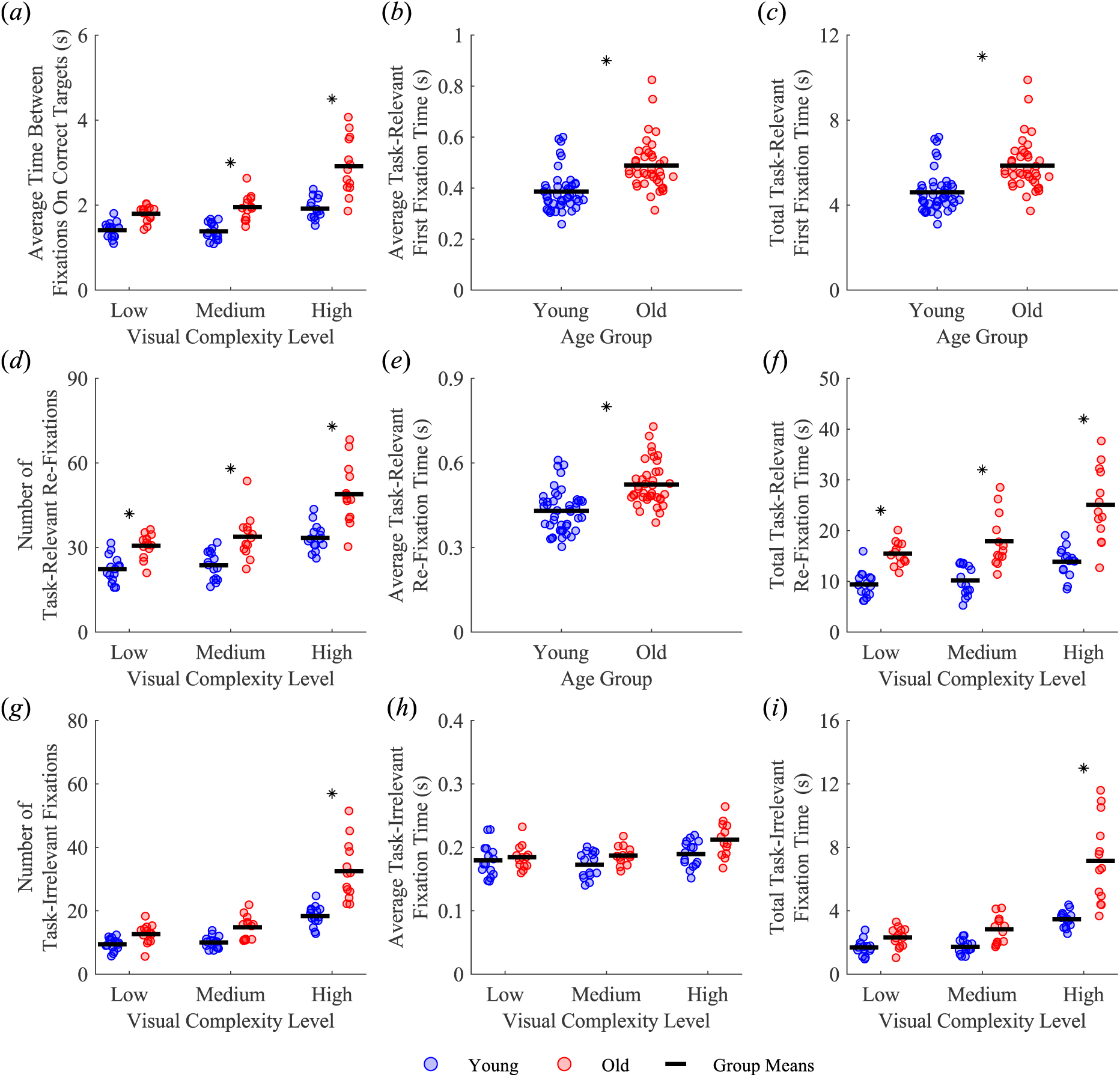
Effect of age group and visual complexity level on gaze behavior. (*a*) Average time between fixations on correct targets, (*b*) average task-relevant first fixation time, (*c*) total task-relevant first fixation time, (*d*) number of task-relevant re-fixations, (*e*) average task-relevant re-fixation time, (*f*) total task-relevant re-fixation time, (*g*) number of task-irrelevant fixations, (*h*) average task-irrelevant fixation time, and (*i*) total task-irrelevant fixation time. *Indicates significant difference between age groups.

**Figure 3.**
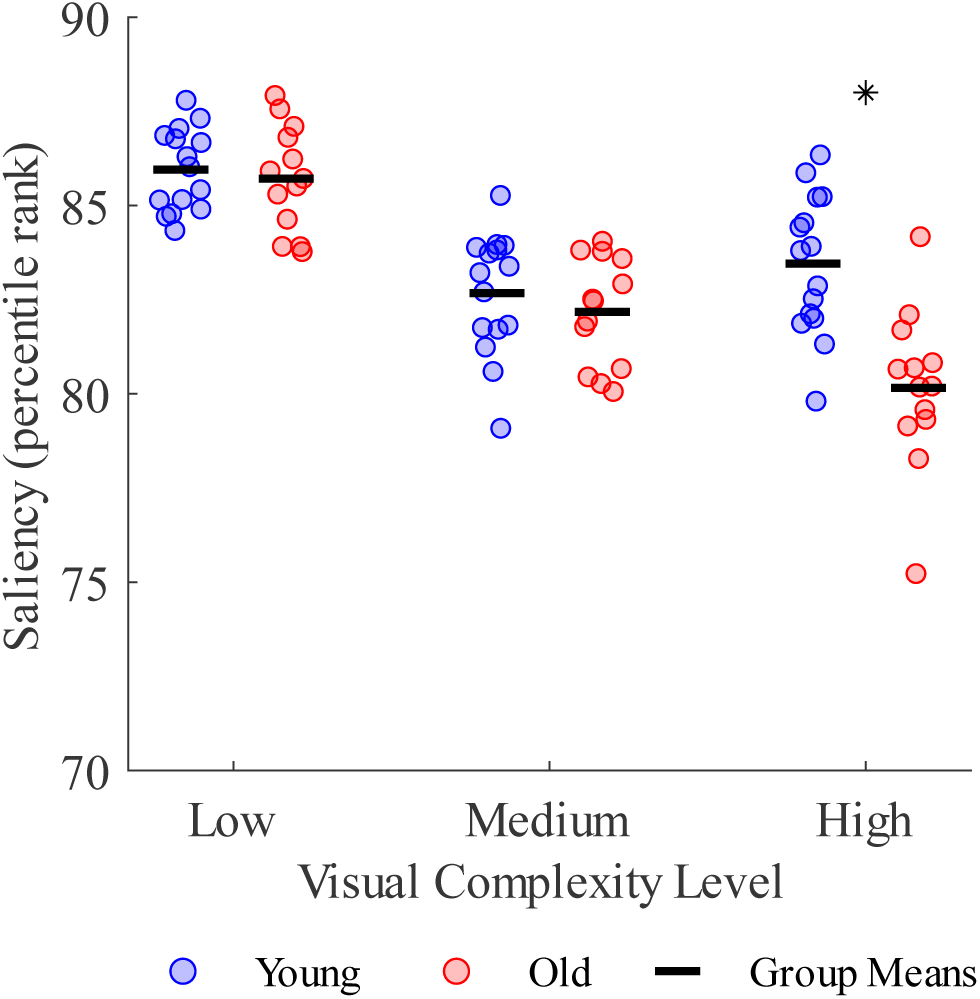
Effect of age group and visual complexity on the saliency of fixated regions. *Indicates significant difference between age groups.

The increase in task completion time in more complex environments may also stem from changes in gaze behavior, such as an increase in fixations on task-relevant versus task-irrelevant objects. We classified fixations into three categories. We considered the first fixation on task-relevant objects as a measure that potentially reflects when participants encoded information about these objects, such as location, in memory. We then identified re-fixations on task-relevant objects as a potential indicator that participants did not recall the location of a given object. Lastly, we measured fixations on task-irrelevant objects as a measure of distractibility. We separately quantified the average duration of each fixation type, total fixation time, and the number of each type of fixation in a trial.

### First fixations on task-relevant objects

We found a main effect of age group on average task-relevant first fixation time (*F*(1,26) = 12.95, *p* = 0.001, Fig. 3b), but there was no significant effect of visual complexity level (*F*(2,52) = 2.56, *p* = 0.087) or interaction between age group and visual complexity level (*F*(2,52) = 1.47, *p* = 0.238). Compared to young adults, older adults spent 0.1 seconds longer on average on their first fixations on task-relevant objects. We also found a main effect of age group on total task-relevant first fixation time (*F*(1,26) = 13.23, *p* = 0.001, Fig. 3c) but not a main effect of visual complexity (*F*(2,52) = 2.85, *p* = 0.067) or an interaction between age group and visual complexity on total (*F*(2,52) = 1.43, *p* = 0.250) task-relevant first fixation times. Compared to young adults, older adults spent 1.3 seconds longer in total on their first fixations on task-relevant objects.

### Re-fixations on task-relevant objects

We calculated the number of re-fixations, the average re-fixation time, and the total re-fixation time on task-relevant objects. We found a main effect of age group (*F*(1,26) = 35.20, *p* < 0.001) and visual complexity (*F*(2,52) = 68.21, *p* < 0.001) on the number of task-relevant re-fixations (Fig. 3d). We also found an interaction effect between age group and visual complexity (*F*(2,52) = 3.91, *p* = 0.026). Compared to young adults, older adults re-fixated task-relevant objects eight more times in the low complexity level (*p* = 0.021), 10 more times in the medium complexity level (*p* = 0.002), and 16 more times in the high complexity level (*p* < 0.001). Additionally, the young adults re-fixated task-relevant objects 11 more times in the high versus low complexity level (*p* < 0.001) and 10 more times in the high versus medium complexity level (*p* < 0.001), while the older adults re-fixated task-relevant objects 18 more times in the high versus low complexity level (*p* < 0.001) and 15 more times in the high versus medium complexity level (*p <* 0.001).

We found a main effect of age group on the average task-relevant re-fixation time (*F*(1,26) = 13.58, *p* = 0.001; Fig. 3e) but did not find a main effect of visual complexity (*F*(2,52) = 1.22, *p* = 0.304) or an interaction effect between age group and visual complexity (*F*(2,52) = 0.01, *p* = 0.994). Compared to the young adults, the older adults spent 0.1 seconds longer on average on their re-fixations on task-relevant objects.

We found main effects of age group (*F*(1,26) = 40.20, *p* < 0.001) and visual complexity (*F*(2,52) = 47.42, *p* < 0.001) on the total task-relevant re-fixation time (Fig. 3f). We also found an interaction effect between age group and visual complexity (*F*(2,52) = 5.97, *p* = 0.005). Compared to the young adults, the older adults spent 6.1 seconds longer in the low complexity level (*p* = 0.005), 7.8 seconds longer in the medium complexity level (*p* < 0.001), and 11.2 seconds longer in the high complexity level (*p* < 0.001) in total on their task-relevant re-fixations. In addition, young adults spent 4.5 seconds longer in the high versus low complexity level (*p* = 0.001) and 3.7 seconds longer in the high versus medium complexity level (*p* = 0.011), while older adults spent 9.6 seconds longer in the high versus low complexity level (*p* < 0.001) and 7.1 seconds longer in the high versus medium complexity level (*p* < 0.001) on their total task-relevant re-fixation times.

### Fixations on task-irrelevant objects

We separately quantified the number of fixations, average fixation time, and total fixation time on task-irrelevant objects. We found main effects of age group (*F*(1,26) = 29.97, *p* < 0.001) and visual complexity (*F*(2,52) = 166.40, *p* < 0.001) on the number of task-irrelevant fixations (Fig. 3g). We also found an interaction effect between age group and visual complexity (*F*(2,52) = 23.16, *p* < 0.001). Compared to the young adults, the older adults fixated on task-irrelevant objects 14 more times in the high complexity level (*p <* 0.001). In addition, the young adults fixated on task-irrelevant objects nine more times in the high versus low complexity level (*p* < 0.001) and eight more times in the high versus medium complexity level (*p* < 0.001) while the older adults fixated on task-irrelevant objects 20 more times in the high versus low complexity level (*p* < 0.001) and 18 more times in the high versus medium complexity level (*p* < 0.001).

We found a main effect of age group (*F*(1,26) = 4.42, *p* = 0.045) on the average task-irrelevant fixation time (Fig. 3h) as the older adults spent 0.01 seconds longer on average on their fixations on task-irrelevant objects. We also found a main effect of visual complexity (*F*(2,52) = 14.57, *p* < 0.001) as all participants spent 0.02 seconds longer in the high versus low complexity (*p* < 0.001) and in the high versus medium complexity (*p* < 0.001) on their average task-irrelevant fixation time. We did not find an interaction effect between age group and visual complexity (*F*(2,52) = 2.16, *p* = 0.125).

We found main effects of age group (*F*(1,26) = 26.36, *p* < 0.001) and visual complexity (*F*(2,52) = 119.33, *p* < 0.001) on the total task-irrelevant fixation time (Fig. 3i). We also found an interaction effect between age group and visual complexity level (*F*(2,52) = 24.17, *p* < 0.001). Compared to the young adults, the older adults spent 3.7 seconds longer on their total task-irrelevant fixation time in the high visual complexity level (*p* < 0.001). In addition, the young adults spent 1.8 seconds longer in the high versus low complexity level (*p* < 0.001) and 1.7 seconds longer in the high versus medium complexity level (*p* < 0.001), while the older adults spent 4.8 seconds longer in the high versus low complexity level (*p* < 0.001) and 4.3 seconds longer in the high versus medium complexity level (*p* < 0.001) in total on their task-irrelevant fixations.

### Saliency of fixated regions

Although both young and older adults increased their fixations on task-irrelevant objects as the visual complexity of the environment increased, these fixations were not towards more salient regions or objects (Fig. 4). We quantified the saliency of the fixated regions on the visual scene using the Itti, Koch, and Neibur model of saliency-based visual attention(52) and found main effects of age group (*F*(1,26) = 10.82, *p* = 0.003) and visual complexity (*F*(2,52) = 60.37, *p* < 0.001). We also found an interaction between age group and visual complexity (*F*(2,52) = 9.27, *p* < 0.001). Young adults fixated more salient regions than older adults in the highest visual complexity level (*p* < 0.001). In addition, young adults fixated more salient regions in the low versus the medium (*p* < 0.001) and high (*p* < 0.001) complexity levels, while older adults fixated more salient regions in the low versus the medium (*p* < 0.001) and high (*p* < 0.001) complexity levels as well as in the medium versus the high (*p* = 0.015) complexity level.

**Figure 4.**
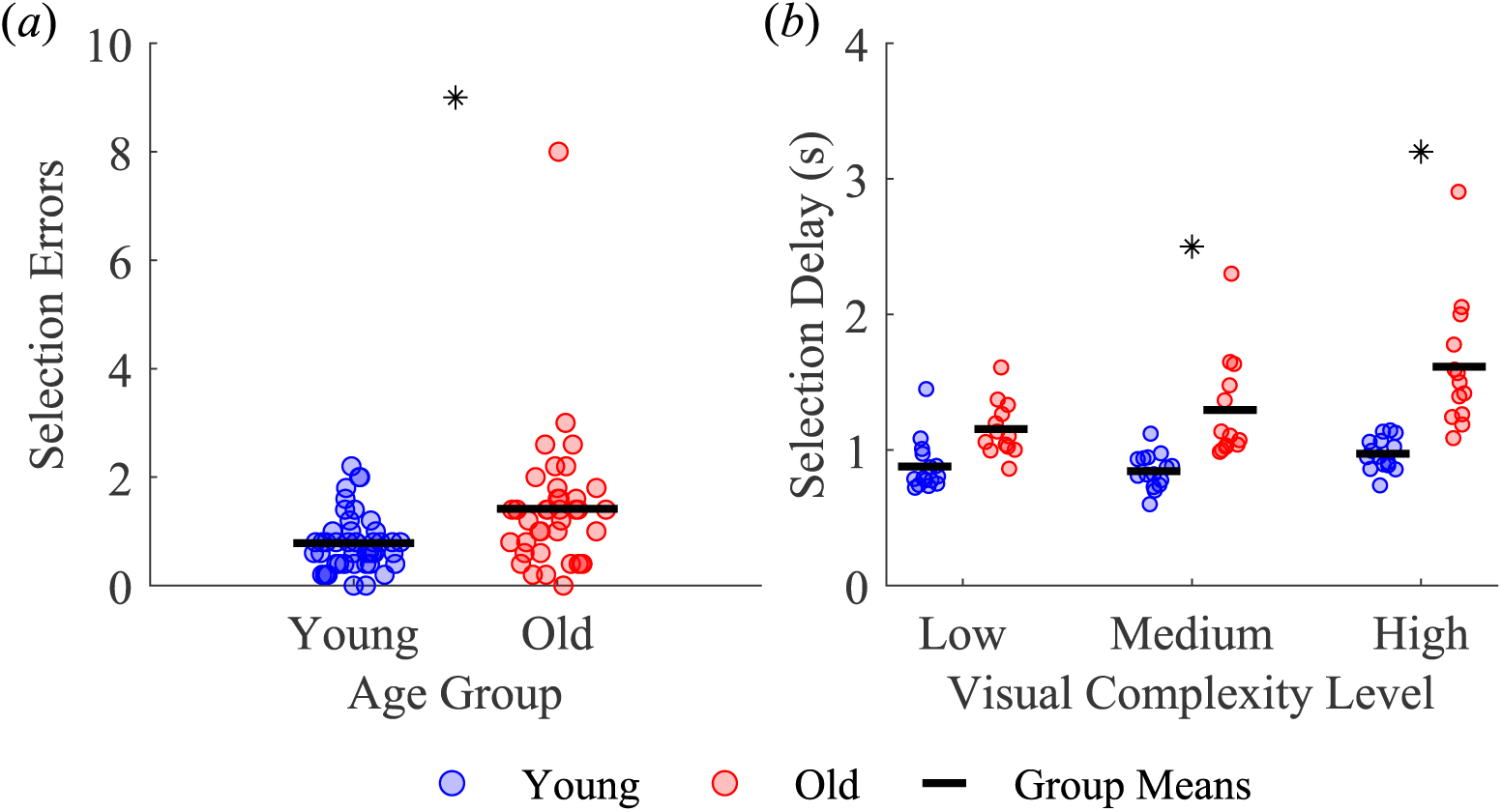
Effect of age group and visual complexity level on (*a*) the number of selection errors and (*b*) the time between fixating and selecting a correct target. *Indicates significant difference between age groups.

### Number of selection errors

Participants may have also taken longer to complete the task with increasing visual complexity due to changes in their target selection strategy. To this end, we quantified target selection errors, defined as selecting an incorrect target or misaiming (laser pointed off-center) when selecting a correct target (Fig. 4a). We found a main effect of age group on the number of selection errors (*F*(1,26) = 6.02, *p* = 0.021), with older adults committing more selection errors than younger adults. We also found a main effect of visual complexity (*F*(2,52) = 3.25, *p* = 0.047) on the number of selection errors; however, post-hoc tests found no differences between visual complexity levels. We did not find an interaction effect between age group and visual complexity (*F*(2,52) = 2.05, *p* = 0.140) on the number of selection errors.

### Selection delay

Another way to investigate possible influences of changes in target selection on task completion time is to quantify selection delay, defined as the average time interval between fixating a correct target and successfully selecting it (Fig. 4b). We found main effects of age group (*F*(1,26) = 34.08, *p* < 0.001) and visual complexity (*F*(2,52) = 11.47, *p* < 0.001) on selection delay. We also found an interaction effect between age group and visual complexity (*F*(2,52) = 4.39, *p* = 0.017) on selection delay. Compared to the young adults, the older adults selected the correct target 0.5 seconds slower in the medium complexity level (*p* = 0.001) and 0.6 seconds slower in the high complexity level (*p* < 0.001). In addition, older adults selected the correct target 0.5 seconds slower in the high versus low complexity level (*p* < 0.001) and 0.3 seconds slower in the high versus medium complexity level (*p* < 0.001).

### Cognitive assessment scores

Age-related differences were observed in several standard cognitive assessments (Table 1). Compared to young adults, older adults had lower global cognition according to their Montreal Cognitive Assessment(41) scores (Wilcoxon rank sum test *p* = 0.044) and poorer short-term memory according to their Corsi Block task(42) results (Wilcoxon rank sum test *p* = 0.001). However, no differences between age groups were found in working memory (two sample t-test *p* = 0.229) as tested using the Backwards Corsi Block task(43) or inhibition (two sample t-test *p* = 0.620) with the Flanker task(44).

**Table 1.** Cognitive assessment scores (Mean ± SD) of young and older adults with the corresponding p-value from their comparisons using two-sample t-tests and Wilcoxon rank sum tests. *Indicates significant differences between age groups.

| Cognitive Assessment | Cognitive Domain | Young Adults (Mean ± SD) | Older Adults (Mean ± SD) | P-value |
| --- | --- | --- | --- | --- |
| Montreal Cognitive Assessment | Global Cognition | 28 ± 2 | 27 ± 3 | 0.0440* |
| Corsi Block Task | Short Term Memory | 6 ± 1 | 5 ± 1 | 0.0014* |
| Backwards Corsi Block Task | Working Memory | 5 ± 1 | 5 ± 2 | 0.229 |
| Flanker Task | Inhibition | 5 ± 70.17 ms | -10.77 ± 95.82 ms | 0.6203 |

### Trail Making Test-B and VR visual search task

As we designed our VR visual search task to mimic several aspects of the paper-based Trail Making Test-B, we were also interested in determining if the outcomes of these assessments were correlated. As such, we fit a multiple linear regression model to determine if performance in our VR visual search task was related to performance in the paper-based Trail Making Test-B (Table 2). In our model, we used Trail Making Test-B completion time as the response and VR task completion time and its interaction with visual complexity level as predictors (*F*(3,80) = 5,61, *p* = 0.002, *Adj. R2* = 0.143). Trail Making Test-B completion time was positively correlated with VR task completion time (*β* = 1.254, *p* < 0.001), but this correlation decreased in the high visual complexity level (*β* = −0.400, *p* = 0.029).

**Table 2.** Results of the multiple linear regression testing for associations between Trail Making Test-B completion time and VR task completion time. Performance in the low visual complexity level was used as the reference.

| Predictor | $\beta$ Estimate | Standard Error | P-value |
| --- | --- | --- | --- |
| VR Task Completion Time | 1.254 | 0.324 | < 0.001 |
| VR Task Completion Time:Medium Visual Complexity Level | -0.080 | 0.171 | 0.641 |
| VR Task Completion Time:High Visual Complexity Level | -0.400 | 0.180 | 0.029 |

### VR visual search task performance and cognitive assessments

Performance in the visual search task could have been influenced by various cognitive domains such as short-term memory, working memory, and inhibitory capacity. As such, we fit a multiple linear regression model to determine if scores on the Montreal Cognitive Assessment, Corsi Block task, Backwards Corsi Block task, and the Flanker task are associated with VR task completion time in the high visual complexity level (Table 3). Additionally, we included age as a predictor to control for potential age-related performance deficits. We specifically chose the high visual complexity level since its range of completion times covered those observed in the lower visual complexity levels. We found significant relationships between VR task performance and cognitive assessment scores (*F*(5,22) = 26.14, *p* < 0.001, *Adj. R2* = 0.82). Specifically, VR task completion times were lower in participants with higher scores on the Montreal Cognitive Assessment (β = −1.42, *p* = 0.005) and Backwards Corsi Block task (β = −2.70, *p* = 0.002). Additionally, VR task completion times increased with age (β = 0.296, *p* < 0.001).

**Table 3.**
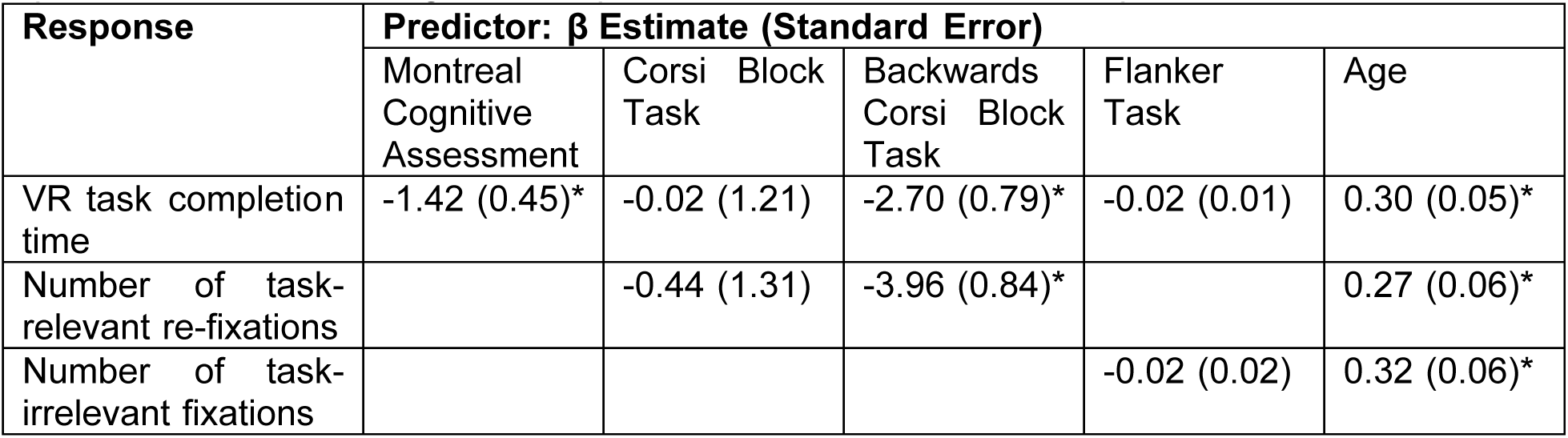
Results of multiple linear regressions testing for associations between performance metrics in the high visual complexity level and cognitive assessment scores. Age was added as a predictor to control for age-related performance deficits. *Indicates p < 0.05.

| Response | Predictor: $\beta$ Estimate (Standard Error) | | | | |
| --- | --- | --- | --- | --- | --- |
|  | Montreal Cognitive Assessment | Corsi Block Task | Backwards Corsi Block Task | Flanker Task | Age |
| VR task completion time | -1.42 (0.45)* | -0.02 (1.21) | -2.70 (0.79)* | -0.02 (0.01) | 0.30 (0.05)* |
| Number of task-relevant re-fixations |  | -0.44 (1.31) | -3.96 (0.84)* |  | 0.27 (0.06)* |
| Number of task-irrelevant fixations |  |  |  | -0.02 (0.02) | 0.32 (0.06)* |

### Gaze behavior and cognitive assessments

Gaze behavior in the task could also act as internal measures of certain cognitive domains. For instance, the number of re-fixations on task-relevant objects could reflect a participant’s short-term and working memory capacity. To test this, we fit a multiple linear regression model to determine if scores in the Corsi Block and Backwards Corsi Block tasks are related to the number of re-fixations on task-relevant objects in the high visual complexity level (Table 3). We included age as a predictor to control for age-related performance deficits and chose to use gaze behavior measurements from the high visual complexity level since its range of values covered those observed in the lower visual complexity levels. Our model explained the data well (*F*(3,24) = 28.99, *p* < 0.001, *Adj. R2* = 0.76) and indicated that the number of re-fixations on task-relevant objects decreased as Backwards Corsi Block score increased (β = −3.96, *p* < 0.001) and increased with age (β = 0.268, *p* < 0.001).

Additionally, the number of fixations on task-irrelevant objects could reflect a participant’s inhibitory capacity. To test this, we fit a separate multiple linear regression model to determine if Flanker task scores are related to the number of fixations on task-irrelevant objects in the high visual complexity level (Table 3). We included age as a predictor to control for age-related performance deficits and chose to use gaze behavior measurements from the high visual complexity level since its range of values covered those observed in the lower visual complexity levels. From this model (*F*(2,25) = 17.58, *p* < 0.001, *Adj. R2* = 0.55), we found that while the number of fixations on task-irrelevant objects increased with age (β = 0.323, *p* < 0.001), it was not associated with scores on the Flanker task (β = −0.019, *p* = 0.218).

## Discussion

We used a VR-based visual search task to determine if increasing the visual complexity of a three-dimensional search environment would lead to changes in performance and if this effect is modulated by age. Additionally, we sought to understand which cognitive domains were associated with performance in VR-based visual search task. For both age groups, increasing the visual complexity of the virtual search environments led to longer task completion times. This effect was due, in part, to the fact that all participants spent more time between fixations on correct targets as they increased their re-fixations on task-relevant and fixations on task-irrelevant objects. We also found that all participants took more time to select a correct target after fixating them in levels with higher visual complexity, which likely contributed to the longer task completion times. Each of these effects was greater in older adults. However, increased completion time did not appear to be influenced by saliency as both groups increasingly fixated on less salient regions as the visual complexity of the environment increased. Lastly, we found that short-term and working memory capacities explained the variability in performance across participants.

As the complexity of the virtual environments increased, all participants spent more time completing the search task because they spent more time searching for correct targets, as demonstrated by the increase in the time interval between fixations on correct targets. These results are consistent with those classically found in simple conjunction search paradigms with simple stimuli composed of arbitrarily selected shapes and colors. When participants searched for stimuli that shared features with distractors, search times increased as the number of distractors in the visual display increased(55). We also found that the effect of visual complexity on search performance was greater in older adults than in young adults. Similar age-related differences in search performance have been demonstrated using simple conjunction search paradigms(56–58) and paradigms using more complex stimuli. For example, when searching for and responding to a star-shaped stimulus in a vehicle’s digital dashboard, older adults demonstrated longer search times as the visual complexity of the display increased than young adults(59). Similar results were observed when comparing search times for specific face configurations(60). It appears then that the influence of visual complexity, age, and their combination on search performance generalizes from two-dimensional displays to more complex, three-dimensional virtual environments.

The increase in suboptimal gaze behavior demonstrated by all participants with increasing visual complexity could have also contributed to the changes in visual search performance. One such suboptimal gaze behavior is the increase in the number of and total time on task-irrelevant fixations with increasing visual complexity, which was found to be greater in older adults. Increased fixations on task-irrelevant objects could be due to a decreased capacity for top-down suppression of irrelevant stimuli with increasing visual complexity of search displays(61). Older adults are also more susceptible to distraction by task-irrelevant information than young adults in various two-dimensional visual-perceptual tasks(21–24,62) and this has been attributed to cognitive impairments, particularly with deficiencies in inhibitory capacity(18,19). However, performance in the Flanker task, a common assessment for inhibitory capacity, was not associated with the number of fixations on task-irrelevant objects and did not differ between age groups in our study. While all participants, particularly older adults, were more prone to directing their attention towards task-irrelevant distractors in higher visual complexity levels in our VR visual search task, it did not seem to be related to their inhibitory capacity.

Another suboptimal gaze behavior observed from all participants at higher visual complexity levels was an increase in the number of and total time on task-relevant re-fixations, with this effect being greater in older than young adults. We also found that scores on the Backwards Corsi Block task, a known test for working memory, were inversely associated with the number of task-relevant re-fixations but did not differ between age groups. It has been previously demonstrated that working memory processes and visual attention are tightly linked. Particularly, increased distraction by task-irrelevant visual stimuli may interfere with encoding task-relevant visual information, ultimately resulting in poor maintenance and retrieval from working memory(25,63). As such, the increased distraction by task-irrelevant objects experienced at higher visual complexity levels may have led to the greater re-fixations observed across all participants.

While all participants became more prone to distraction by task-irrelevant stimuli as visual complexity increased, this effect was not driven by the saliency of as the participants increasingly fixated regions that have lower saliency values as the visual complexity of the virtual environments increased. While prior studies showed that susceptibility to distraction during visual perception tasks is greater in the presence of salient visual stimuli, particularly in older adults(21,23,62), this effect appears to be subdued in more complex scenes as people weigh prior knowledge of or the gist of the scene more than saliency(13–15). In our task, participants may have formed a gist of each visual complexity level during practice and the initial five trials in a visual complexity level. Specifically, they would have learned that the search environments are modeled after farms, the animals are located naturally at ground level, and the letters are arbitrarily placed in the sky. The emphasis on prior knowledge and gist instead of bottom-up saliency when directing gaze and visual attention in realistic images(64–67) appears to generalize to three-dimensional virtual environments. As such, the ability of visual stimuli to automatically capture visual attention may not be contingent upon the stimuli being highly salient.

Proper allocation of visual attention is important for completing visual search tasks and appears to be associated with cognition. Performance in our task was related to global cognition and working memory capacity, as those who scored higher in the Montreal Cognitive Assessment and the Backwards Corsi block task demonstrated shorter task completion times. These results echo the importance of working memory capacity on visual attention(68–70), which has been demonstrated in simple selective attention(71) and visual search(60) tasks. Beyond memory, inhibitory capacity is another cognitive domain that influences the allocation of visual attention and the successful completion of visual search tasks. As previously discussed, inhibitory capacity allows for the proper allocation of visual attention by selecting for and processing information relevant to the task while suppressing those that are irrelevant. However, we found no relationship between inhibitory capacity, which was tested using the Flanker task, and VR task completion times. Visual search performance and the relevant gaze behavior that may influence it appear to be influenced by an individual’s general cognitive capacity, with working memory playing a more prominent role, but not inhibition.

The increase in task completion times at higher visual complexity levels can be explained not only by the time required to search for the correct targets, as exhibited by the calculated fixation times, but also by changes that influence the speed with which targets are selected, such as by moving slower. We quantified selection delay as the time between fixating and selecting a correct target, which is influenced by a combination of the time it takes to move the laser pointer to the correct target and successfully select it. All participants demonstrated longer selection delay as the visual complexity of the search environments increased, with this effect being greater in older adults. These effects on selection delay could have been due to participants spending more time during information processing to determine whether the target is correct relative to the sequence. However, we did not find an effect of visual complexity on average fixation times on task-relevant objects. Longer selection delays at higher visual complexity levels could also have been caused by a general movement slowing as participants traded off speed for accuracy(72,73). This effect seems to be supported by the lack of changes in the number of selection errors with increasing visual complexity for all participants. However, the selection errors were greater in older adults than young adults, contradicting previously demonstrated parity between the two groups resulting from older adults using movement slowing as a compensatory mechanism to maintain accuracy(72,73).

It is important to consider that our study contains limitations that may have influenced our results and their interpretability. While the task design improves on prior visual search tasks on static two-dimensional realistic images, using letters in the search sequence may have diminished the ecological validity of our task. Additionally, the search targets in our tasks were related (letters of the alphabet and visual representation of animals whose names started with those letters) such that participants only needed to keep track of a single set in their working memory. In contrast, the paper-based Trail Making Test-B uses two different, unrelated lists (letters of the alphabet and numbers), requiring working memory to keep track of two sets. The low adjusted R2 we found when determining the relationship between performance in our VR task and the paper-based Trail Making Test-B also supports this limitation. Future work may benefit from creating a VR-based visual search task that uses only search targets that can be found in the real world and fully incorporates all aspects of the paper-based Trail Making Test-B. While all of our participants have normal or corrected-to-normal vision, we were not able to assess the visual acuity and contrast sensitivity of our participants, which may influence a person’s perception of saliency since it is quantified using the low-level visual features of a scene and ultimately the results in our study. Future work should include assessments that target various visual domains. Finally, the lack of a relationship between inhibitory capacity and the various performance metrics used may be due to our sample of older adults being cognitively high functioning and the assessment used not targeting the type of inhibition present during visual search. Future work would benefit from including participants with a broader range of cognitive assessment scores and using a more comprehensive cognitive assessment battery targeting more cognitive domains.

In conclusion, time to complete the VR visual search task increased with the visual complexity of the environment as participants increased time on re-fixating task-relevant objects and fixating task-irrelevant objects. This negative effect was greater in older adults and is associated with differences in cognitive capacity, particularly in working memory. Increased susceptibility to distraction is potentially problematic as suboptimal allocation of visual attention may lead to difficulties completing everyday tasks and can potentially lead to injuries. For instance, young and older adults who show greater anxiety stemming from fear of falling while walking demonstrate greater susceptibility to distraction by threatening stimuli, which may negatively affect their ability to adapt their gait for safe locomotion(74–76). These results highlight the importance of designing cognitive assessments that use tasks that closely simulate the visual complexity and dynamicity in the real world. In addition, our study also emphasizes the need for a more comprehensive analysis of task performance beyond completion time to accurately assess visuomotor behavior, particularly in older adults.

## Data Accessibility

The processed data and the code used to generate the figures and statistics are available on the Open Science Framework at https://osf.io/r9q3k/.

## Author Contributions

IJL and JMF conceived and designed the experiment. IJL and AK developed the VR-based visual search task. IJL collected the data. IJL and JMF analyzed the data and interpreted the results. IJL and JMF drafted, edited, and revised the manuscript. IJL, AK, and JMF approved the final version of the manuscript.

## Ethical Statement

All study procedures have been reviewed and approved by University of Southern California’s Institutional Review Board (UP-22-00317). All participants provided their written informed consent before the start of the experiment and all aspects of study conformed to the principles expressed in the Declaration of Helsinki of 1975, as revised in 2013.

## References

1. Land MF, Hayhoe M. In what ways do eye movements contribute to everyday activities? Vision Res. 2001;41(25–26):3559–65.

2. Land MF. Eye movements and the control of actions in everyday life. Prog Retin Eye Res. 2006;25:296–324.

3. Hayhoe M, Ballard D. Eye movements in natural behavior. Trends Cogn Neurosci. 2005;9(4):188–94.

4. Hayhoe MM, Rothkopf CA. Vision in the natural world. Wiley Interdiscip Rev Cogn Sci. 2011 Mar 1;2(2):158–66.

5. Patla AE. Understanding the roles of vision in the control of human locomotion. Gait Posture [Internet]. 1997;5(1):54–69. Available from: http://www.sciencedirect.com/science/article/pii/S0966636296011095

6. Marigold DS, Patla AE. Gaze fixation patterns for negotiating complex ground terrain. Neuroscience. 2007 Jan 5;144(1):302–13.

7. Matthis JS, Yates JL, Hayhoe MM. Gaze and the Control of Foot Placement When Walking in Natural Terrain. Curr Biol. 2018 Apr 23;28(8):1224–1233.e5.

8. Chapman GJ, Hollands MA. Evidence for a link between changes to gaze behaviour and risk of falling in older adults during adaptive locomotion. Gait Posture. 2006;24(3):288–94.

9. Chapman GJ, Hollands MA. Evidence that older adult fallers prioritise the planning of future stepping actions over the accurate execution of ongoing steps during complex locomotor tasks. Gait Posture. 2007;26(1):59–67.

10. Young WR, Hollands MA. Can telling older adults where to look reduce falls? Evidence for a causal link between inappropriate visual sampling and suboptimal stepping performance. Exp Brain Res. 2010 Jul;204(1):103–13.

11. Itti L, Koch C. Computational modelling of visual attention. Nat Rev Neurosci. 2001;2(3):194– 203.

12. Tatler BW, Hayhoe MM, Land MF, Ballard DH. Eye guidance in natural vision: Reinterpreting salience. J Vis. 2011;11(5).

13. Wolfe JM, Võ MLH, Evans KK, Greene MR. Visual search in scenes involves selective and nonselective pathways. Trends Cogn Sci. 2011 Feb 1;15(2):77–84.

14. Wolfe JM. Visual search. Curr Biol. 2010 Apr 27;20(8):R346–9.

15. Henderson JM. Human gaze control during real-world scene perception. Trends Cogn Sci. 2003;7(11):498–504.

16. Mather M. Aging and cognition. WIREs Cogn Sci. 2010;1(3):346–62.

17. Harada CN, Natelson Love MC, Triebel K. Normal Cognitive Aging. Clin Geriatr Med. 2013 Nov;29(4):737–52.

18. Dempster FN. The rise and fall of the inhibitory mechanism: Toward a unified theory of cognitive development and aging. Dev Rev. 1992 Mar 1;12(1):45–75.

19. Lustig C, Hasher L, Zacks RT. Inhibitory deficit theory: Recent developments in a “new view.” In: Gorfein DS, Macleod CM, editors. Inhibition in cognition. American Psychological Association; 2007. p. 145–62.

20. Gazzaley A, Mark D’esposito A. Top-Down Modulation and Normal Aging. Ann N Y Acad Sci. 2007;1097:67–83.

21. Kramer AF, Hahn S, Irwin DE, Theeuwes J. Age differences in the control of looking behavior: Do You Know Where Your Eyes Have Been? Psychol Sci. 2000;11(3):210–7.

22. Schmitz TW, Cheng FHT, De Rosa E. Failing to ignore: paradoxical neural effects of perceptual load on early attentional selection in normal aging. J Neurosci Off J Soc Neurosci. 2010 Nov 3;30(44):14750–8.

23. Schmitz TW, Dixon ML, Anderson AK, De Rosa E. Distinguishing attentional gain and tuning in young and older adults. Neurobiol Aging. 2014 Nov 1;35(11):2514–25.

24. Mertes C, Wascher E, Schneider D. Compliance instead of flexibility? On age-related differences in cognitive control during visual search. Neurobiol Aging. 2017 May 1;53:169– 80.

25. Gazzaley A, Clapp W, Kelley J, McEvoy K, Knight RT, D’Esposito M. Age-related top-down suppression deficit in the early stages of cortical visual memory processing. Proc Natl Acad Sci. 2008 Sep 2;105(35):13122–6.

26. Reitan RM. Validity of the Trail Making Test as an indicator of organic brain damage. Percept Mot Skills. 1958;8(7):271.

27. Sánchez-Cubillo I, Periáñez JA, Adrover-Roig D, Rodríguez-Sánchez JM, Ríos-Lago M, Tirapu J, et al. Construct validity of the Trail Making Test: Role of task-switching, working memory, inhibition/interference control, and visuomotor abilities. J Int Neuropsychol Soc. 2009;15(3):438–50.

28. Salthouse TA. What cognitive abilities are involved in trail-making performance? Intelligence. 2011 Jul;39(4):222.

29. Llinàs-Reglà J, Vilalta-Franch J, López-Pousa S, Calvó-Perxas L, Torrents Rodas D, Garre-Olmo J. The Trail Making Test. Assessment. 2017 Mar 1;24(2):183–96.

30. Cangoz B, Karakoc E, Selekler K. Trail Making Test: Normative data for Turkish elderly population by age, sex and education. J Neurol Sci. 2009 Aug 15;283(1):73–8.

31. Hashimoto R, Meguro K, Lee E, Kasai M, Ishii H, Yamaguchi S. Effect of age and education on the Trail Making Test and determination of normative data for Japanese elderly people: The Tajiri Project. Psychiatry Clin Neurosci. 2006;60(4):422–8.

32. Specka M, Weimar C, Stang A, Jöckel KH, Scherbaum N, Hoffmann SS, et al. Trail Making Test Normative Data for the German Older Population. Arch Clin Neuropsychol. 2022 Feb 1;37(1):186–98.

33. Suzuki H, Sakuma N, Kobayashi M, Ogawa S, Inagaki H, Edahiro A, et al. Normative Data of the Trail Making Test Among Urban Community-Dwelling Older Adults in Japan. Front Aging Neurosci. 2022 May 25;14.

34. Tombaugh TN. Trail Making Test A and B: Normative data stratified by age and education. Arch Clin Neuropsychol. 2004 Mar 1;19(2):203–14.

35. Plotnik M, Ben-Gal O, Doniger GM, Gottlieb A, Bahat Y, Cohen M, et al. Multimodal immersive trail making-virtual reality paradigm to study cognitive-motor interactions. J NeuroEngineering Rehabil. 2021 Dec 1;18(1):1–16.

36. Recker L, Poth CH. Test–retest reliability of eye tracking measures in a computerized Trail Making Test. J Vis. 2023 Aug 22;23(8):15.

37. Wölwer W, Falkai P, Streit M, Gaebel W. Trait Characteristic of Impaired Visuomotor Integration during Trail-Making Test B Performance in Schizophrenia. Neuropsychobiology. 2003 Sep 26;48(2):59–67.

38. Singh T, Perry CM, Fritz SL, Fridriksson J, Herter TM. Eye Movements Interfere With Limb Motor Control in Stroke Survivors. Neurorehabil Neural Repair. 2018 Aug 1;32(8):724–34.

39. Adlakha G, Singh S, Patil A, Nuthalapati K, Khandve P, Bhattacharyya P, et al. Development of a virtual reality assessment of visuospatial function and oculomotor control. In: 2021 IEEE Conference on Virtual Reality and 3D User Interfaces Abstracts and Workshops (VRW). Institute of Electrical and Electronics Engineers Inc.; 2021. p. 753–4.

40. Kreidler SM, Muller KE, Grunwald GK, Ringham BM, Coker-Dukowitz ZT, Sakhadeo UR, et al. GLIMMPSE: Online Power Computation for Linear Models with and without a Baseline Covariate. J Stat Softw. 2013 Sep;54(10):i10.

41. Nasreddine ZS, Phillips NA, Bédirian V, Charbonneau S, Whitehead V, Collin I, et al. The Montreal Cognitive Assessment, MoCA: A Brief Screening Tool For Mild Cognitive Impairment. J Am Geriatr Soc. 2005 Apr 1;53(4):695–9.

42. Corsi PM. Memory and the medial temporal region of the brain. [Montreal]: McGill University; 1972.

43. Isaacs EB, Vargha-Khadem F. Differential course of development of spatial and verbal memory span: A normative study. Br J Dev Psychol. 1989 Nov;7(4):377–80.

44. Eriksen BA, Eriksen CW. Effects of noise letters upon the identification of a target letter in a nonsearch task. Percept Psychophys. 1974 Jan;16(1):143–9.

45. Stoet G. PsyToolkit: A software package for programming psychological experiments using Linux. Behav Res Methods. 2010 Nov;42(4):1096–104.

46. Stoet G. PsyToolkit: A Novel Web-Based Method for Running Online Questionnaires and Reaction-Time Experiments. Teach Psychol. 2017 Jan 1;44(1):24–31.

47. Rosenholtz R, Li Y, Nakano L. Measuring visual clutter. J Vis. 2007 Jan 2;7(2):17–17.

48. Matin E. Saccadic suppression: a review and an analysis. Psychol Bull. 1974 Dec;81(12):899–917.

49. Rayner K. Eye movements and attention in reading, scene perception, and visual search. Q J Exp Psychol 2006. 2009;62(8):1457–506.

50. Salvucci DD, Goldberg JH. Identifying Fixations and Saccades in Eye-Tracking Protocols. In: Proceedings of the EyeTracking Research and Applications Symposium. New York: Association for Computing Machinery; 2000. p. 71–8.

51. Tobii Pro Lab Gaze Filter [Internet]. 2023 [cited 2024 Jul 15]. Tobii Pro Lab Gaze Filter. Available from: https://connect.tobii.com

52. Itti L, Koch C, Niebur E. A model of saliency-based visual attention for rapid scene analysis. IEEE Trans Pattern Anal Mach Intell. 1998;20(11):1254–9.

53. Le Meur O, Baccino T. Methods for comparing scanpaths and saliency maps: Strengths and weaknesses. Behav Res Methods. 2013;45(1):251–66.

54. Kretch KS, Adolph KE. Active vision in passive locomotion: Real-world free viewing in infants and adults. Dev Sci. 2015 Sep 1;18(5):736–50.

55. Treisman AM, Gelade G. A feature-integration theory of attention. Cognit Psychol. 1980;12(1):97–136.

56. Folk CL, Lincourt AE. The effects of age on guided conjunction search. Exp Aging Res. 1996 Jan 1;22(1):99–118.

57. Hommel B, Li KZH, Li SC. Visual search across the life span. Dev Psychol. 2004 Jul;40(4):545–58.

58. Müller-Oehring EM, Schulte T, Rohlfing T, Pfefferbaum A, Sullivan EV. Visual Search and the Aging Brain: Discerning the Effects of Age-related Brain Volume Shrinkage on Alertness, Feature Binding, and Attentional Control. Neuropsychology. 2013;27(1):48.

59. Lee SC, Kim YW, Ji YG. Effects of visual complexity of in-vehicle information display: Age-related differences in visual search task in the driving context. Appl Ergon. 2019 Nov 1;81:102888.

60. Aziz JR, Good SR, Klein RM, Eskes GA. Role of aging and working memory in performance on a naturalistic visual search task. Cortex. 2021;136:28–40.

61. Bonacci LM, Bressler S, Kwasa JAC, Noyce AL, Shinn-Cunningham BG. Effects of Visual Scene Complexity on Neural Signatures of Spatial Attention. Front Hum Neurosci. 2020 Mar 24;14:91.

62. Tsvetanov KA, Mevorach C, Allen H, Humphreys GW. Age-related differences in selection by visual saliency. Atten Percept Psychophys. 2013;75(7):1382–94.

63. Gazzaley A, Nobre AC. Top-down modulation: bridging selective attention and working memory. Trends Cogn Sci. 2012 Feb 1;16(2):129–35.

64. Henderson JM, Malcolm GL, Schandl C. Searching in the dark: Cognitive relevance drives attention in real-world scenes. Psychon Bull Rev. 2009 Oct 1;16(5):850–6.

65. Henderson JM, Hayes TR. Meaning guides attention in real-world scene images: Evidence from eye movements and meaning maps. J Vis. 2018;18(6).

66. Peacock CE, Hayes TR, Henderson JM. Meaning guides attention during scene viewing, even when it is irrelevant. Atten Percept Psychophys. 2019;81(1):20–34.

67. Peacock CE, Hayes TR, Henderson JM. The role of meaning in attentional guidance during free viewing of real-world scenes. Acta Psychol (Amst). 2019 Jul 1;198.

68. Hayhoe MM. Vision and Action. Annu Rev Vis Sci. 2017;3:389–413.

69. Madden DJ. Aging and Visual Attention. Curr Dir Psychol Sci. 2007 Apr;16(2):70.

70. Diamond A. Executive Functions. Annu Rev Psychol. 2013 Jan 2;64:135–68.

71. Shipstead Z, Harrison TL, Engle RW. Working memory capacity and visual attention: Top-down and bottom-up guidance. Q J Exp Psychol. 2012 Mar;65(3):401–7.

72. Zhang L, Yang J, Inai Y, Huang Q, Wu J. Effects of aging on pointing movements under restricted visual feedback conditions. Hum Mov Sci [Internet]. 2015 Apr 1 [cited 2025 Feb 20];40:1–13. Available from: https://www.sciencedirect.com/science/article/pii/S016794571400205X

73. Lamb DG, Correa LN, Seider TR, Mosquera DM, Rodriguez JA, Salazar L, et al. The aging brain: Movement speed and spatial control. Brain Cogn [Internet]. 2016 Nov 1 [cited 2023 Oct 27];109:105–11. Available from: https://www.sciencedirect.com/science/article/pii/S0278262616301634

74. Ellmers TJ, Young WR. The influence of anxiety and attentional focus on visual search during adaptive gait. J Exp Psychol Hum Percept Perform. 2019 Jun 1;45(6):697–714.

75. Ellmers TJ, Cocks AJ, Kal EC, Young WR. Conscious movement processing, fall-related anxiety, and the visuomotor control of locomotion in Older Adults. J Gerontol - Ser B Psychol Sci Soc Sci. 2020 Nov 1;75(9):1911–20.

76. Young WR, Wing AM, Hollands MA. Influences of state anxiety on gaze behavior and stepping accuracy in older adults during adaptive locomotion. J Gerontol - Ser B Psychol Sci Soc Sci. 2012;67 B(1):43–51.

